# Ectopic assembly of an auxin efflux control machinery shifts developmental trajectories

**DOI:** 10.1101/2023.09.16.558043

**Authors:** Ana Cecilia Aliaga Fandino, Adriana Jelinkova, Petra Marhava, Jan Petrasek, Christian S. Hardtke

**Affiliations:** Department of Plant Molecular Biology, University of Lausanne, CH-1015 Lausanne, Switzerland; Institute of Experimental Botany, Czech Academy of Sciences, 165 02 Prague, Czech Republic

**Keywords:** Arabidopsis, auxin, AGC kinase, xylem, differentiation

## Abstract

Polar auxin transport in the Arabidopsis root tip maintains high auxin levels around the stem cell niche that gradually decrease in dividing cells but increase again once they transition towards differentiation. Protophloem differentiates earlier than other proximal tissues and employs a unique auxin ‘canalization’ machinery that is thought to balance auxin efflux with retention. It consists of a proposed activator of PIN auxin efflux carriers, the AGC kinase PAX; its inhibitor, BRX; and PIP5K enzymes, which promote polar PAX and BRX localization. Because of dynamic PAX-BRX-PIP5K interplay, the net cellular output of this machinery remains unclear. Here we deciphered the dosage-sensitive regulatory interactions between PAX, BRX and PIP5K by their ectopic expression in developing xylem vessels. The data suggest that the dominant collective output of the PAX-BRX-PIP5K module is a localized reduction in PIN abundance. This requires PAX-stimulated clathrin-mediated PIN endocytosis by site-specific phosphorylation, which distinguishes PAX from other AGC kinases. Importantly, ectopic assembly of the PAX-BRX-PIP5K module is sufficient to cause cellular auxin retention and affects root growth vigor by accelerating the trajectory of xylem vessel development. Our data thus provide direct evidence that local manipulation of auxin efflux alters the timing of cellular differentiation in the root.

## Introduction

The phytohormone auxin regulates plant development as well as adaptive responses by modulating growth patterns. Auxin action depends both on context and concentration, and is determined by an interplay of auxin biosynthesis, transport and signaling (Adamowski and Friml, 2015; Lavy and Estelle, 2016; Zhao, 2018). Long distance transport occurs in bulk through the plant vascular system, whereas short distance, cell-to-cell transport depends on dedicated plasma-membrane-integral auxin carriers (Morris and Kadir, 1972; Teale et al., 2006; Adamowski and Friml, 2015). They comprise the auxin influx facilitator AUX1 and its homologs, and the PIN-FORMED (PIN) auxin efflux carriers. The latter are chiefly responsible for creating the high local auxin concentrations that are observed in the growth apices of plants, the meristems (Blilou et al., 2005). Auxin maxima are associated with the formation of new, lateral organs, but are also required to maintain the meristems themselves. For example, the auxin maximum at the tip of *Arabidopsis thaliana* (Arabidopsis) root meristems is essential for the establishment and maintenance of the stem cell niche (SCN) (Sabatini et al., 1999). It is created by coordinated, generally rootward polar subcellular localization of PIN proteins in the stele and ground tissue, and reinforced by an “inverse fountain” of auxin recycling mediated by shootward-pointing PINs in the columella and epidermis (Grieneisen et al., 2007). The auxin maximum thus is the peak of an auxin gradient that determines the activity of transcriptional regulators, which in turn specify the different tissue layers and time their proliferation and differentiation (Mahonen et al., 2014).

PIN protein localization is a dynamic process that involves endocytic recycling and associated regulatory mechanisms. For example, phosphorylation of the cytoplasmic hydrophilic loop by the AGC kinase PINOID (PID) can induce PIN re-localization as well as depolarization (Friml et al., 2004; Weller et al., 2017; Wang et al., 2023). Other AGC family kinases such as D6 PROTEIN KINASE (D6PK) also target phosphosites in the hydrophilic loop of PINs but thereby activate PIN-mediated auxin efflux from the cytoplasm into the apoplast (Willige et al., 2013; Barbosa et al., 2014; Zourelidou et al., 2014). PIN localization also depends on the low abundant plasma membrane phosphoinositide phosphatidylinositol-4,5-bisphosphate [PI(4,5)P2], which affects clathrin-mediated PIN endocytosis (Ischebeck et al., 2013; Tejos et al., 2014). PI(4,5)P2 is produced from the more abundant phosphatidylinositol-4-phosphate (PI4P) by PHOSPHATIDYLINOSITOL-4-PHOSPHATE-5-KINASE (PIP5K) enzymes, like the redundant PIP5K1 and PIP5K2 in the Arabidopsis root.

The early vascular tissues of the root meristem, the protophloem and protoxylem, are formed inside the stele in a diarch pattern, wherein two protophloem poles are flanking an axis of metaxylem vessels that is delimited by protoxylem cell files on both sides (Fig. S1A). Molecular markers highlight both developing xylem vessels and protophloem sieve elements (the conducting cells of the protophloem) as sites of auxin accumulation (Bishopp et al., 2011; Marhava et al., 2018), which is thought to promote their differentiation (Bishopp et al., 2011; Vaughan-Hirsch et al., 2018; Moret et al., 2020; von der Mark et al., 2022; Wang et al., 2023). For example, in *pip5k1 pip5k2* double mutants, frequent differentiation failures of xylem vessel precursors are associated with low auxin levels and can be partially rescued by induction of local auxin production (von der Mark et al., 2022). PIP5K1/2 are predominantly found in association with the plasma membrane but are also present in the nucleus (Gerth et al., 2017; Watari et al., 2022), and both subcellular localizations are required for normal xylem vessel development (von der Mark et al., 2022).

*pip5k1 pip5k2* double mutants also display severe protophloem sieve element differentiation failures (Marhava et al., 2020). In developing sieve elements, PIP5K1/2 display a strongly polar, rootward plasma membrane association, which is conferred by interaction with a ‘molecular rheostat’ composed of BREVIS RADIX (BRX) and the AGC kinase PROTEIN KINASE ASSOCIATED WITH BRX (PAX) (Marhava et al., 2020; Wang et al., 2023). Together, the three proteins form an interdependent self-reinforcing polarity module that regulates auxin efflux and responds itself to auxin. Briefly, the current model suggests that BRX inhibits PAX-mediated auxin efflux activation at low cellular auxin levels, while the recruitment of PIP5K reinforces PAX localization because PI(4,5)P2 promotes PAX polarity (Barbosa et al., 2016). Upon rising auxin levels, PAX activity is potentiated by 3-phosphoinositide-dependent protein kinase (PDK)-mediated phosphorylation (Marhava et al., 2018; Xiao and Offringa, 2020). Subsequently, PAX activates auxin efflux by phosphorylating PINs as well as BRX, which is consequently displaced from the plasma membrane (Marhava et al., 2018; Koh et al., 2021; Wang et al., 2023). Because BRX is required for efficient PIP5K recruitment, and because cellular auxin levels drop due to efflux, the system is eventually reset (Aliaga Fandino and Hardtke, 2022; Wang et al., 2023). The ensuing dynamic equilibrium coordinates auxin flux between adjacent cells to prevent the emergence of fate bistability and leads to auxin canalization in the developing sieve element file (Moret et al., 2020; Aliaga Fandino and Hardtke, 2022).

One cellular output of the self-reinforcing rheostat system is a subcellular PIN pattern that is specific for developing protophloem sieve elements (Marhava et al., 2020). That is, co-localized PIP5K, PAX and BRX association with the center of the rootward plasma membrane in a ‘muffin’ domain creates a local minimum of PIN abundance which therefore appears as a complementary ‘donut’ pattern (Fig. S1B and C). Markers suggest that this central minimum is possibly created by clathrin-mediated PIN endocytosis (Marhava et al., 2020; Wang et al., 2023). In *pax*, *brx* or *pip5k1 pip5k2* mutants, PIN abundance is increased and displays the even ‘pancake’ distribution throughout the plasma membrane as observed in other cell files (Marhava et al., 2020; Wang et al., 2023). Ultimately, it is BRX-tampered PAX activity that creates the PIN minimum, whereas PIP5K is mainly required to promote PAX polarity in antagonism to sieve element-specific CLAVATA3/EMBRYO SURROUNDING REGION-RELATED 45 (CLE45) peptide signaling through its receptor BARELY ANY MERISTEM 3 (BAM3) (Wang et al., 2023).

Since the protophloem is essential for root meristem maintenance and growth (Anne and Hardtke, 2017), the sieve element differentiation failures in *pax*, *brx* or *pip5k1 pip5k2* mutants are accompanied by a short root phenotype (Marhava et al., 2018; Marhava et al., 2020). Although the observed fate bistability in these loss-of-function backgrounds supports the idea that auxin accumulation is required for sieve element formation (Marhava et al., 2018; Moret et al., 2020), the systemic effects of perturbed protophloem development also obscure the potential role of post-SCN auxin increase in timing the transition to differentiation. Moreover, PAX-mediated PIN control is required for root growth vigor even in the absence of visible protophloem differentiation defects (Wang et al., 2023), raising the question how the tradeoff between PIN activation and PIN abundance plays out. Here we built on the knowledge that auxin accumulation is required for xylem vessel differentiation (von der Mark et al., 2022), and that a spatio-temporal shift in xylem differentiation does not necessarily affect overall root growth (Ramachandran et al., 2021) to address these issues directly, via a gain-of-function approach in an ectopic context.

## Results

### PAX-mediated PIN1 phosphorylation promotes PIN1 endocytosis

The interdependence of BRX-PAX-PIP5K module assembly and polarity was previously demonstrated through protophloem sieve element (PPSE)-specific induction of corresponding CITRINE fusion proteins under control of an estradiol-inducible *COTYLEDON VASCULAR PATTERN 2* promoter *(CVP2^XVE^)* in reciprocal mutant backgrounds (Wang et al., 2023). Here we used such transgenic lines to determine the impact of the individual components on subcellular PIN patterning, thereby also exploiting the fact that the fusion proteins are by comparison over-expressed upon prolonged induction (Figure 1) (Fig. 1A). As previously reported (Wang et al., 2023), PPSE-specific induction of PID, employed as a control, led to a nearly comprehensive PIN1 depolarization (Fig. 1B). PAX induction in *pax* mutant background initially restored the PIN1 ‘donut-to-pancake’ ratio to Columbia-0 (Col-0) wildtype levels. Upon prolonged PAX induction, ‘donuts’ became sharper with a widened PIN1 minimum and even more frequent than in wildtype (Fig. 1A and B). By contrast, induced BRX overexpression in *brx* mutant background eventually led to a strong increase in the ‘pancake’ pattern (Fig. 1A and B). By comparison, prolonged PIP5K1 induction in phenotypically wildtype *pip5k2* single mutant background at best slightly increased the ‘pancake’ frequency (Fig. 1A and B). Collectively, these findings corroborate that PAX is responsible for subcellular PIN ‘donut’ patterning and that BRX inhibits PAX activity (Marhava et al., 2020; Wang et al., 2023).

**Figure 1.**
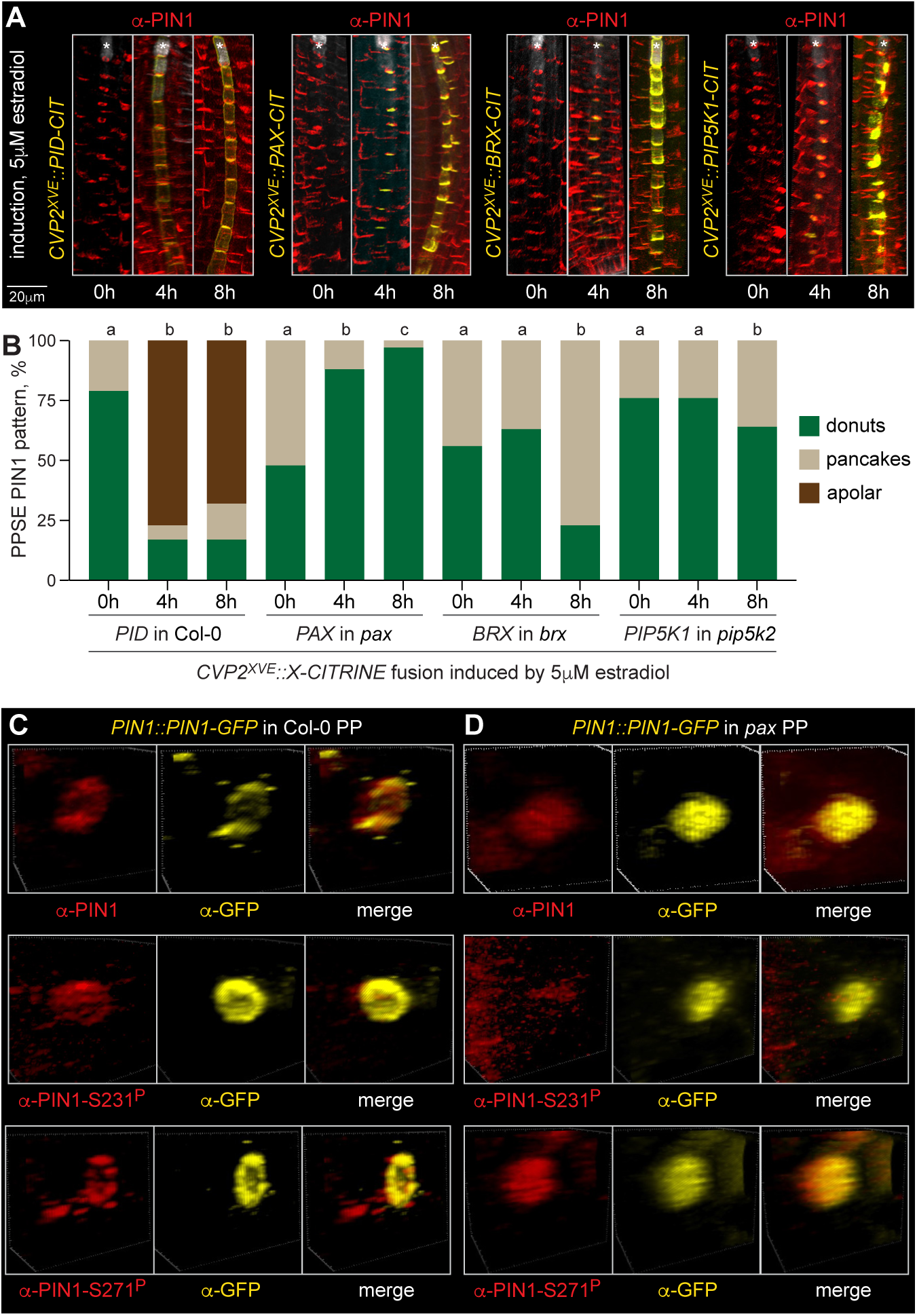
PAX targets a specific PIN1 phosphosite in developing protophloem sieve elements (PPSEs). (A) Simultaneous detection of PIN1 (anti-PINl antibody, red fluorescence) and CITRINE fusion proteins (anti-GFP antibody, yellow fluorescence) by immunostaining. Transgenic plants expressing the indicated fusion proteins under control of the PPSE-specific estradiol-inducible *COTYLEDON VASCULAR PATTERN 2* promoter *(CVP2^XVE^)* were transferred onto estradiol media and monitored at indicated timepoints. Asterisks highlight the PPSE cell file (calcofluor white staining, grey fluorescence). (B) Quantification of the subcellular PIN1 pattern in developing PPSEs, corresponding to (A). n=76-217 PPSEs per time point; statistically significant differences (lower case letters) were determined by Chi square test, p<0.001. (C-D) Simultaneous detection of transgenic PIN1-GFP fusion protein (anti-GFP antibody, yellow fluorescence) with either anti-PINl, or S231^p^-phosphosite-specific anti-PINl, or S271^p^-phosphosite-specific anti-PINl antibodies (red fluorescence) by immunostaining in Columbia-0 (Col-0) wildtype (C) or *pax* mutant (D) background.

AGC kinases phosphorylate several target sites in the hydrophilic loop of PIN proteins, whose combinatorial read-out determines both PIN activity and polarity (Bassukas et al., 2022). For PID, three major target sites in PIN1 have been described, which could be detected with phosphosite-specific anti-PIN1 antibodies (Weller et al., 2017). Two such antibodies were available to us, and we performed immunostainings that corroborated earlier results (Marhava et al., 2018). That is, S271 phosphorylation could still be readily detected in developing PPSEs of *pax* mutants, whereas S231 phosphorylation was essentially absent (Fig. 1C and D). Induction of PAX in *pax* mutant background restored S231 phosphorylation (Fig. S2A), suggesting that S231 is a valid PAX target *in vivo*. Because PAX induction triggers the appearance of pronounced PIN1 ‘donut’ patterns and because PAX kinase activity is required for PIN1 patterning (Wang et al., 2023), the data suggest that S231 phosphorylation promotes PIN1 turnover.

Reduced PAX kinase activity also attenuates PIN recycling as revealed by a reduction of PIN1-containing brefeldin-A (BFA) bodies upon BFA treatment (Wang et al., 2023). Together with the observation that DYNAMIN-RELATED PROTEIN 1 A (DRP1A), a promoter of clathrin-mediated endocytosis (Fujimoto et al., 2010), colocalizes with the BRX-PAX-PIP5K ‘muffin’ domain specifically in developing PPSEs (Dettmer et al., 2014; Marhava et al., 2020), this suggests that the PIN1 minimum may be created by increased local PIN endocytosis. Extended live imaging of developing PPSEs in transgenics expressing an RFP-tagged PIN1 protein simultaneously with either a CITRINE-tagged PAX or BRX protein indeed captured highly dynamic localization of all three proteins, with internalized PIN1 vesicles seemingly originating from the ‘muffin’ domain (Figure 2) (Fig. 2A, Fig. S2B). To test the involvement of clathrin-mediated endocytosis in creating the PIN minimum directly, we made transgenic lines for PPSE-specific estradiol-inducible expression of the dominant-negative endocytosis blockers AUXILIN-LIKE 2 (Adamowski et al., 2018) and C-HUB (Kitakura et al., 2011). Indeed, both AUXILIN-LIKE 2 and C-HUB induction triggered a gradual decrease in the PIN1 ‘donut’ pattern and a corresponding increase in the ‘pancake’ pattern (Fig. 2B-D). In summary, our data support the notion that PAX-mediated PIN1 phosphorylation triggers PIN1 endocytosis to create a local PIN1 minimum.

**Figure 2.**
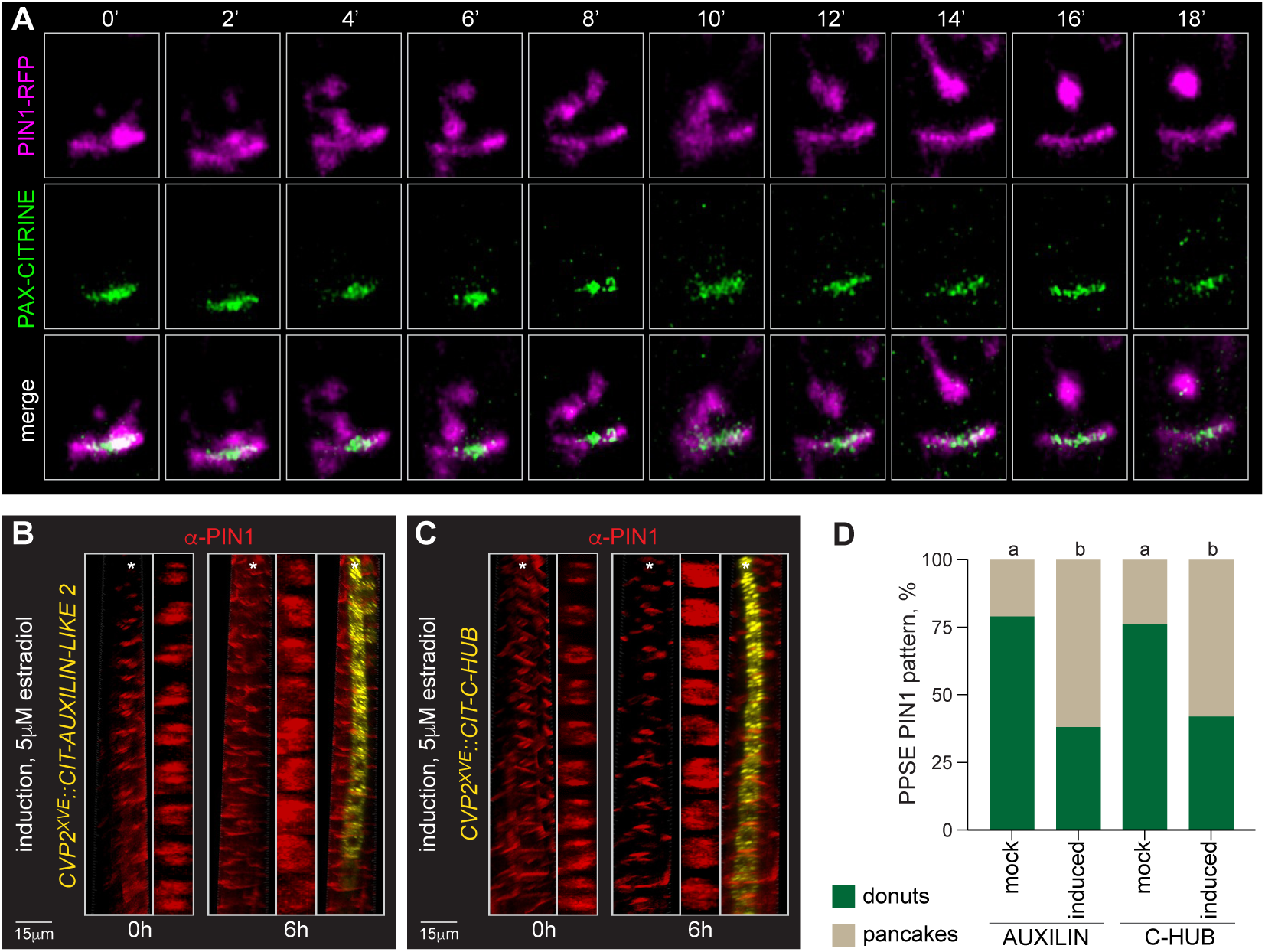
The central minimum in developing PPSEs reflects enhanced PIN1 endocytosis. (A) Time course of PIN1-RFP (magenta fluorescence) and PAX-CITRINE (green fluorescence) fusion protein dynamics at the rootward plasma membrane of a developing PPSE, capturing PIN1-RFP internalization from the center. (B-C) Simultaneous im mu nostaining of PIN 1 (anti-PIN 1 antibody, red fluorescence) and CITRINE fusions (anti-GFP antibody, yellow fluorescence) with dominant inhibitors of clathrin-mediated endocytosis. Transgenic plants expressing either AUXILIN-LIKE 2 (B) or C-HUB (C) fusion protein under control of the *CVP2^XVE^* promoter were monitored before and after transfer onto estradiol media. 3D reconstructions of PIN1 and corresponding top-down views on the rootward end of individual vessels are shown aside merged views with the induced effectors. Asterisks highlight the PPSE cell file (calcofluor white staining, grey fluorescence). (D) Quantification of the subcellular PIN1 pattern in developing PPSEs, corresponding to (B) and (C). n=140-153 PPSEs per time point; statistically significant differences (lower case letters) were determined by Fisher’s exact test, p<0.0001.

### PAX also patterns subcellular PIN1 distribution in developing xylem vessels

PAX is most prominently expressed in developing PPSEs (Figure 3) (Fig. 3A) but also in the xylem axis (Marhava et al., 2018), with a relatively stronger expression in the protoxylem than in the metaxylem (Fig. 3B). Expression of a PAX-CITRINE fusion protein under control of the PPSE-specific *BRX* promoter (Fig. S2C) rescued both protophloem differentiation defects as well as diminished root growth of *pax* mutants however (Marhava et al., 2018) (Fig. S2D and E), suggesting that PAX activity in the developing xylem is not essential for root meristem growth vigor. Nevertheless, simultaneous immunolocalization of PAX fusion protein and PIN1 revealed that similar to developing PPSEs, PAX is also localized in a central ‘muffin’ domain in developing xylem vessels (Fig. 3C and D). Moreover, in these cells PIN1 also frequently displayed a less defined ‘donut’ pattern with a smaller yet recognizable minimum (Fig. 3E) that was not observed in other cell files where PAX was undetectable. In the developing xylem of *pax* mutants, the abundance of this pattern was substantially reduced (Fig. 3F-H). Moreover, live imaging of PIN1 fusion protein signal in the plasma membrane of individual cells over time revealed that PIN1 turnover in developing metaxylem vessels is nearly as dynamic as in developing PPSEs (Fig. S3A). Unlike in neighboring procambial cell files that displayed overall lower PIN1 turnover, PIN1 dynamics were strongly reduced in developing PPSEs of *pax* mutants, and in tendency possibly also in metaxylem vessels (Fig. S3A). In summary, our results suggest that even relatively low amounts of PAX can generate a weak yet recognizable PIN1 ‘donut’ pattern in developing xylem vessels.

**Figure 3.**
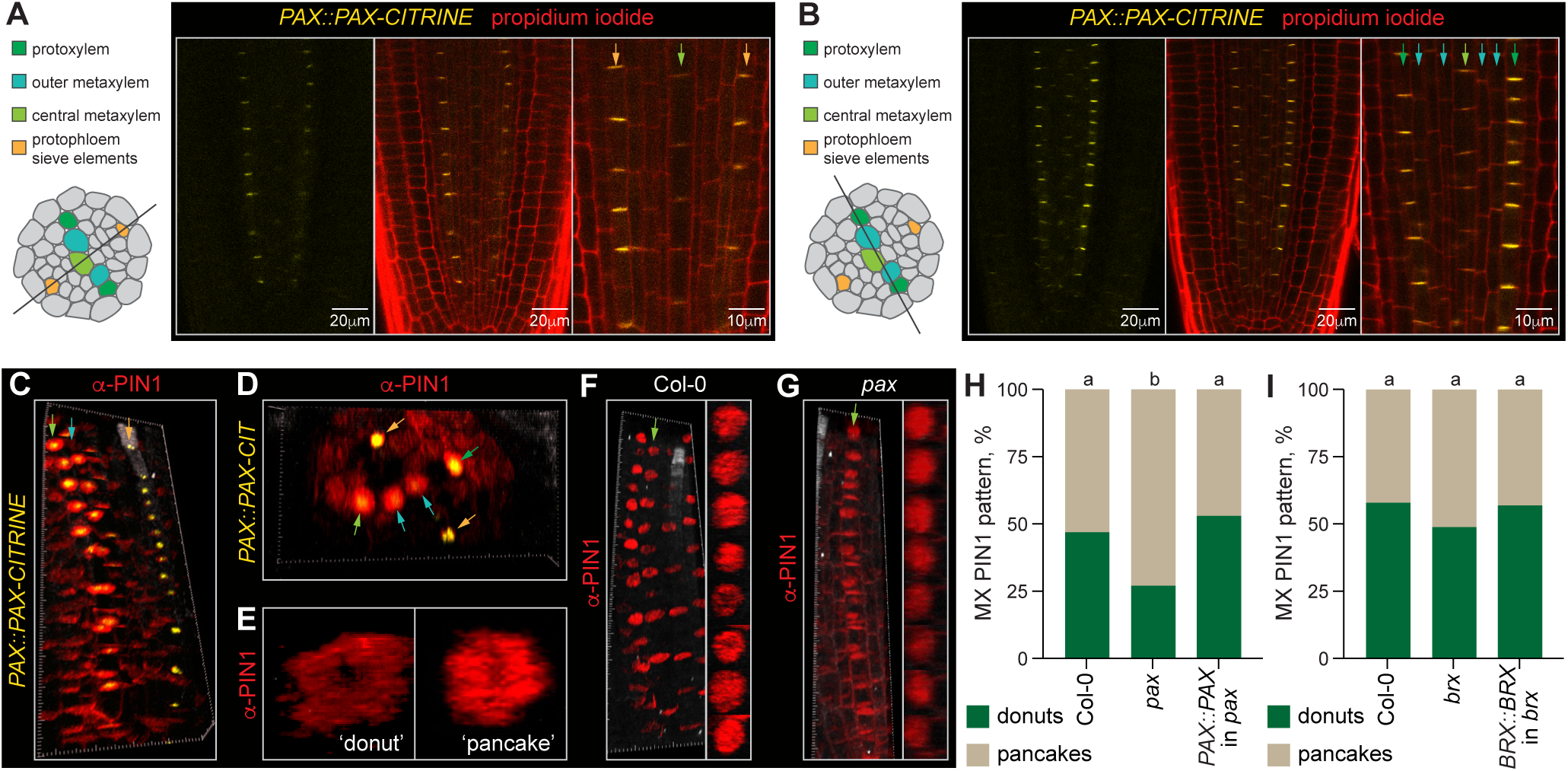
PAX expression in the xylem and corresponding subcellular PIN1 pattern. (A-B) Confocal live imaging of PAX-CITRINE fusion protein (yellow fluorescence, left panels) expressed under control of its native promoter in *pox* mutant background, and merged with propidium iodide cell wall staining (red fluorescence, center panels). Longitudinal optical sections through the protophloem (A) and xylem axis (B) planes are shown. Vascular cell types indicated by arrows in the magnified images (right panels) are color-coded with reference to the schematic overviews. (C-D) Simultaneous detection of PIN1 (anti-PINl antibody, red fluorescence) and PAX-CITRINE fusion protein (anti-GFP antibody, yellow fluorescence) by immunostaining, shown in longitudinal (C) and horizontal (D) view 3D reconstructions. (E) Examples of PIN1 ‘donut’ and ‘pancake’ subcellular patterning in developing metaxylem vessels, detected by anti-PINl antibody staining. (F-G) Detection of PIN1 by anti-PINl antibody staining (red fluorescence) in developing metaxylem vessels, showing 3D reconstructions (left panels) and corresponding top-down views on the rootward end of individual vessels (right panels). (H-l) Quantification of the subcellular PIN1 pattern in developing metaxylem (MX) vessels in indicated genotypes. n=323-483 MX vessels; statistically significant differences (lower case letters) were determined by Fisher’s exact test, p=0.0052.

### BRX antagonizes PAX-mediated PIN1 patterning

Unlike PAX, BRX is not detectable outside developing PPSEs (Marhava et al., 2018) and consistently, *brx* mutants did not show a change in the abundance of xylem vessel PIN1 ‘donuts’ (Fig. 3I). Likewise, components of the CLE45 signaling pathway, which interferes with PAX polarity and thereby PIN1 patterning in developing PPSEs (Wang et al., 2023) (Fig. S3B), are not expressed in the xylem (Kang and Hardtke, 2016; Breda et al., 2019), and consistently CLE45 treatments did not impact subcellular PIN1 patterning in developing metaxylem vessels (Fig. S3C). Thus, the developing xylem is an ideal tissue to probe the functioning of the BRX-PAX-PIP5K module and its cellular impact.

To ectopically assemble the module, we first expressed a BRX-CITRINE fusion protein under control of the *PAX* promoter. As expected, this construct complemented the *brx* PPSE differentiation defects (Fig. S4A) and root growth phenotype (Figure 4) (Fig. 4A). Although *BRX* expression in the *PAX* domain thus had no detrimental effect *per se*, it interfered with the root growth rescue normally conferred by a *PAX::PAX-CITRINE* transgene in *pax* single mutants (Fig. 4B) (Fig. S4B). This was observed in trans-heterozygous *brx +/-pax +/-* background (Fig. 4B), confirming a gain-of-function effect. Importantly, compared to endogenous PAX protein, PAX-CITRINE fusion protein was always expressed at higher levels in developing xylem (Fig. 4C), possibly because of transgene concatenation. However, by itself this did not result in more frequent or more accentuated PIN1 minima (Fig. 3H). In contrast, additional BRX fusion protein expression in the xylem disrupted subcellular PIN1 patterning and led to an increase in the ‘pancake’ configuration (Fig. 4D-F), which is again consistent with BRX being an inhibitor of PAX activity (Marhava et al., 2018). In summary, we found that ectopic expression of BRX in developing xylem vessels interfered with root elongation and correlated with an increase in the PIN1 ‘pancake’ pattern when combined with elevated PAX fusion protein levels.

**Figure 4.**
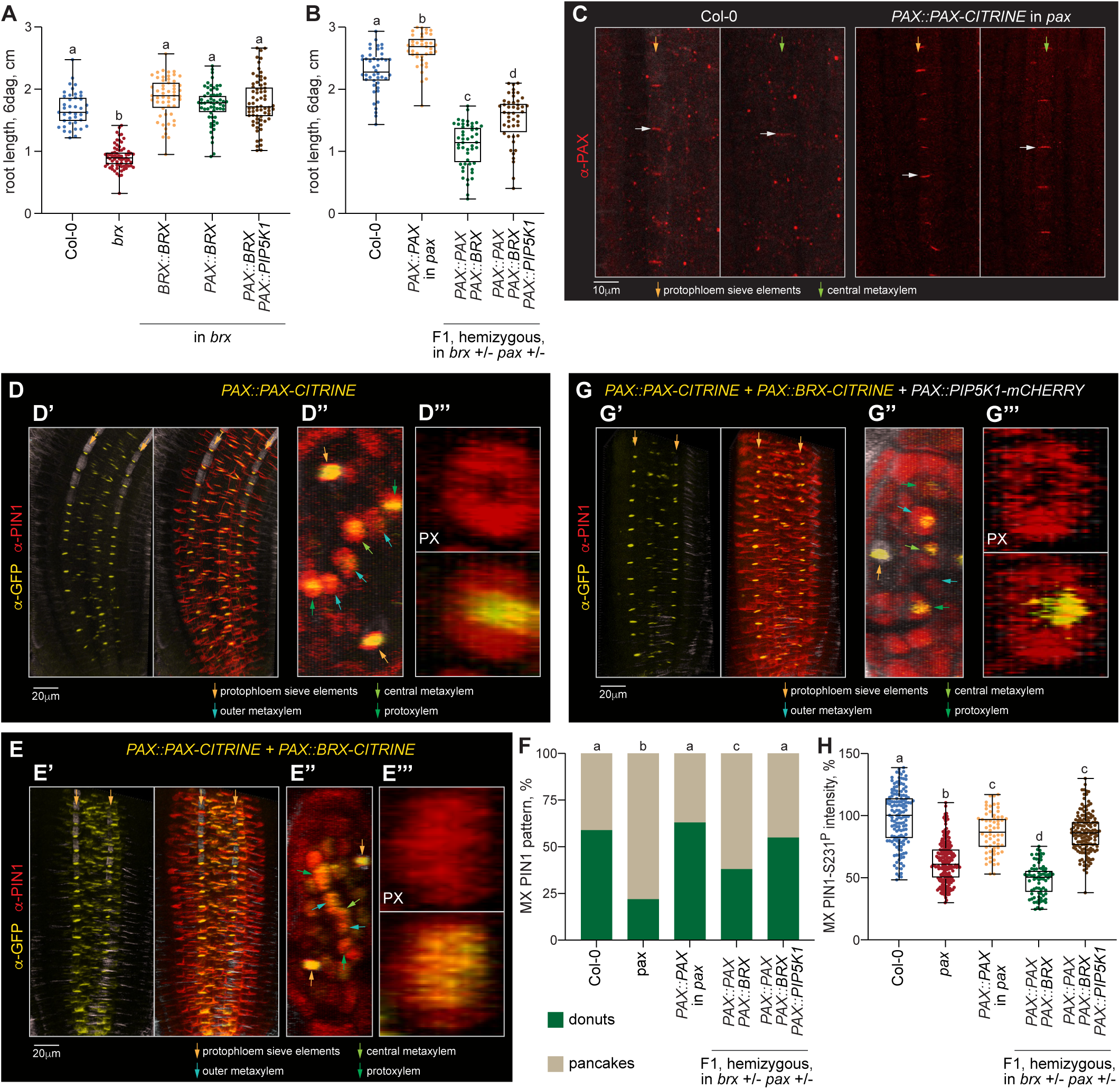
Ectopic expression of the PAX-BRX-PIP5K1 module in developing xylem vessels affects subcellular PIN1 patterning. (A-B) Primary root length of indicated genotypes. Transgenic PAX and BRX proteins were expressed as CITRINE fusions, PIP5K1 as an mCHERRY fusion. n=41-68 roots (A) and n=39-47 roots (B); statistically significant differences were determined by ordinary one-way ANOVA, p<0.0001 in (A) and (B). (C) Detection of native PAX in Col-0 wildtype or transgenic PAX-CITRINE fusion protein in *pox* mutant background by anti-PAX antibody staining (red fluorescence) in developing PPSEs (left panels) or MX vessels (right panels). Note the higher expression level of transgenic fusion protein (e.g. white arrows) as compared to endogenous PAX. (D-E) Simultaneous detection of PIN1 (anti-PINl antibody, red fluorescence) and indicated CITRINE fusion proteins (anti-GFP antibody, yellow fluorescence) by immunostaining, shown in longitudinal (’) and horizontal (“) overview, and top-down view in individual protoxylem (PX) vessels (’“) (3D reconstructions). (F) Quantification of the subcellular PIN1 pattern in developing MX vessels in indicated genotypes. n=142-182 MX vessels; statistically significant differences (lower case letters) were determined by Fisher’s exact test, p≤0.0202. (G) As in D-E. (H) Relative signal intensity of S23l^p^-specific PIN1 immunostaining in developing MX vessels of indicated genotypes. n=62-206 MX vessels; statistically significant differences (lower case letters) were determined by ordinary one-way ANOVA, p≤0.0007. Box plots display 2nd and 3rd quartiles and the median, bars indicate maximum and minimum.

### PIP5K1 dampens PAX inhibition by BRX

Similar to PAX, PIP5K1 and PIP5K2 are both expressed in the developing xylem vasculature albeit at barely detectable levels (von der Mark et al., 2022) (Fig. S4C and D). Unlike PAX however, PIP5K localization in the xylem is not polar, and moreover not only plasma-membrane-associated but also nuclear PIP5K1 is required for proper xylem differentiation (von der Mark et al., 2022). The pronounced polar localization of PIP5K1 in developing PPSEs largely depends on the presence of BRX (Marhava et al., 2020; Wang et al., 2023), and indeed PIP5K1-mCHERRY fusion protein that was expressed in the *PAX* expression domain simultaneously with BRX-CITRINE fusion protein displayed a markedly polar enrichment in the xylem (Fig. S4E) (but not without BRX-CITRINE (Fig. S4F)). Moreover, the PIP5K1 dosage increase partially reversed the negative impact of BRX dosage increase on PAX activity, as indicated by partially recovered root growth (Fig. 4B) and largely restored PIN1 patterning (Fig. 4F and G).

Because PDK1 is expressed in the xylem (Xiao and Offringa, 2020) (Fig. S4G), we also monitored auxin-induced plasma-membrane-dissociation of ectopically expressed BRX. In the *brx* mutant background, BRX-CITRINE fusion protein expressed under control of the *PAX* promoter displayed the expected response in PPSEs but not in metaxylem (Fig. S4H). However, the response in PPSEs was ‘sharpened’ by a PAX dosage increase and could then also be observed in the metaxylem (Fig. S4H). Similar to the other characteristics we had quantified, additional PIP5K1 dampened this response (Fig. S4H). Finally, consistent with our observations, PIN1 S231 phosphorylation was strongly reduced when BRX was introduced into the xylem, but recovered by additional PIP5K1 (Fig. 4H). In summary, these findings reiterate the importance of S231 phosphorylation for the creation of the PIN1 minimum, the positive effect of PIP5K1 on PAX activity, and the intricate quantitative and dosage-sensitive relation between the three module components (Aliaga Fandino and Hardtke, 2022; Wang et al., 2023).

### Ectopic assembly of the PAX-BRX-PIP5K1 module changes the developmental trajectory of xylem cells

The PAX-BRX-PIP5K module has an important role in guiding the transition of developing PPSEs towards differentiation (Marhava et al., 2018; Marhava et al., 2020; Moret et al., 2020), and we thus sought to investigate whether the observed gain-of-function effects were associated with altered developmental trajectories of the xylem. The secondary cell wall pattern is an easily scorable morphological indicator of xylem vessel differentiation status and also distinguishes protoxylem vessels with their reticulated pattern from metaxylem vessels with their pitted pattern (Ramachandran et al., 2021).

First, we monitored xylem vessel patterns in the post-meristematic region of roots, between 5 to 7 mm from the tip. As expected (Graeff and Hardtke, 2021), in this area, protoxylem vessels were always differentiated whereas metaxylem vessels showed some variation between genotypes (Figure 5) (Fig. 5A). In wildtype, we always observed two differentiated protoxylem vessels, two differentiated outer metaxylem vessels, and with very few exceptions an undifferentiated central metaxylem vessel (Fig. 5A and B). In *pax* mutants, the central metaxylem had often already differentiated and occasionally an additional xylem cell file was observed (Fig. 5A and C), and this phenotype could be complemented by a *PAX::PAX-CITRINE* transgene (Fig. 5A and D). Thus, *PAX* loss-of-function may confer a weak xylem phenotype, which however may also simply be related to its short root phenotype because similar aberrations were observed in *brx* mutants (Fig. S5A). Addition of a *PAX::BRX-CITRINE* transgene to the *PAX::PAX-CITRINE* transgene led to more frequent changes in xylem cell file number (Fig. 5A and E) and was accentuated by a *PAX::PIP5K1-CITRINE* transgene (Fig. 5A and F). Compared to wildtype, in the latter triple transgenic we also frequently observed differentiated metaxylem (Fig. 5F).

**Figure 5.**
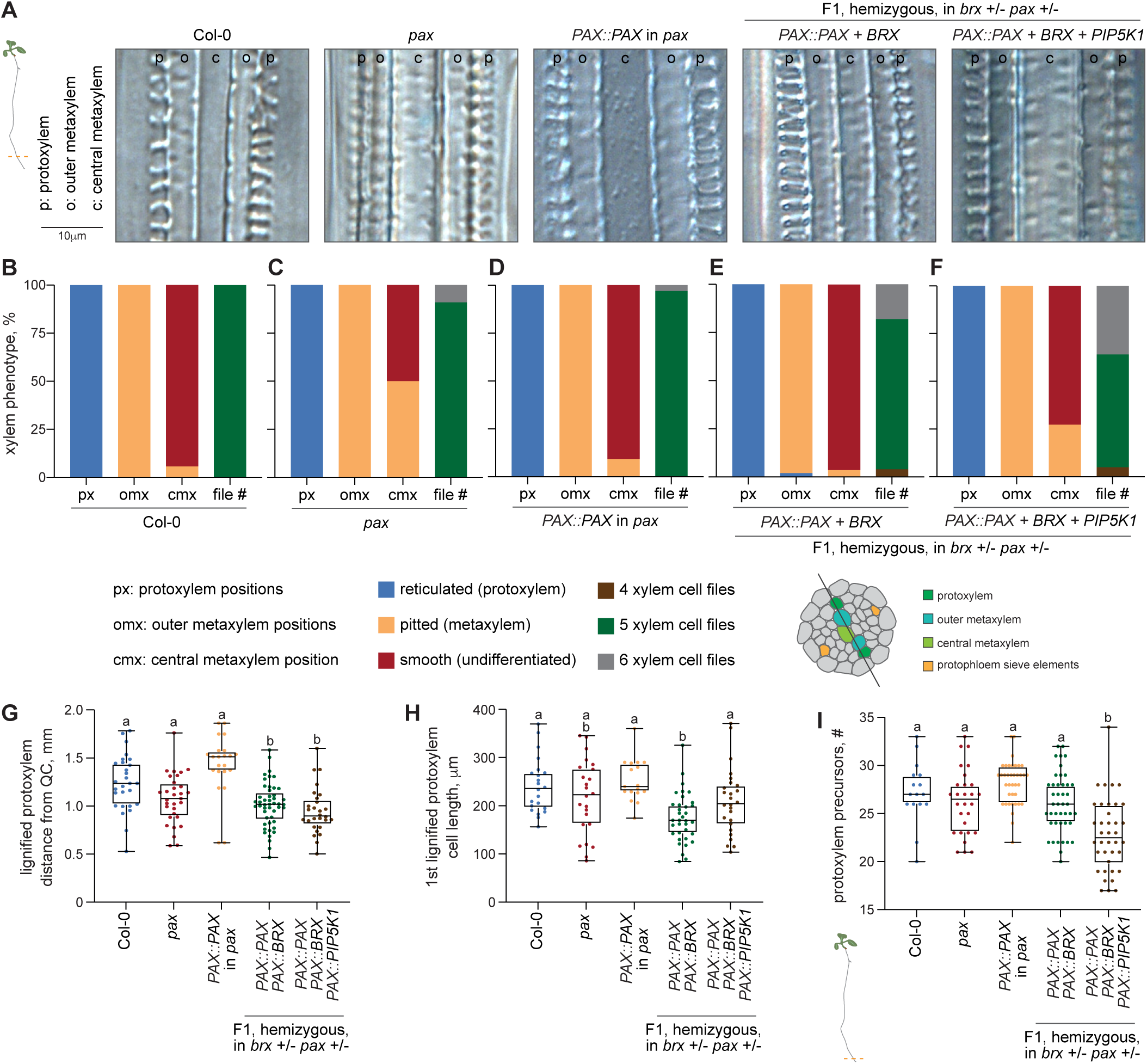
Ectopic assembly of the PAX-BRX-PIP5K1 module affects the trajectory of xylem development. (A-F) Differential interference contrast light microscopy example images of the xylem axis in the indicated genotypes, taken 5-7 mm above the root tip (A), and quantification of corresponding differentiation status per vessel type and genotype (B-F). n=22-35 roots. (G) Distance of the first lignified PX vessels from the quiescent center (QC) in the indicated genotypes. n=22-48 roots; statistically significant differences (lower case letters) were determined by ordinary one-way ANOVA, p≤0.0027. (H) Length of the first lignified PX vessels in the indicated genotypes. n=17-35 PX vessels; statistically significant differences (lower case letters) were determined by ordinary one-way ANOVA, p≤0.OOlO. (I) Number of undifferentiated vessel precursors in PX cell files until the first lignified PX vessel in the indicated genotypes, counted from the QC. n=16-43 cell files; statistically significant differences (lower case letters) were determined by ordinary one-way ANOVA, p=0.0003. Box plots display 2nd and 3rd quartiles and the median, bars indicate maximum and minimum.

Next, we inspected protoxylem differentiation, which occurs closer to the root tip and can be traced continuously from the SCN (Graeff and Hardtke, 2021; Ramachandran et al., 2021). In wildtype, *pax* mutants or complemented *pax* mutants we did not observe a statistically significant difference in the onset of protoxylem differentiation with respect to the distance from the SCN (Fig. 5G). However, protoxylem vessels appeared to differentiate closer to the SCN both when BRX, or BRX and PIP5K1 were combined with increased PAX dosage (Fig. 5G). However, unlike in the triple transgenic situation (PAX + BRX

+ PIP5K1), in the double transgenics (PAX + BRX) we also observed shorter protoxylem cells (Fig. 5H). Finally, we found significantly fewer protoxylem precursor cells in lines expressing the entire PAX-BRX-PIP5K1 module but not in the other genotypes (Fig. 5I). By contrast, no differences were observed in the number of PPSE precursors (Fig. S5B). Thus, ectopic expression of BRX in the xylem together with a PAX dosage increase resulted in overall shorter cells but did not accelerate the trajectory of protoxylem differentiation, whereas ectopic expression of the entire PAX-BRX-PIP5K1 module did. In summary, our data indicate that manipulation of PAX activity in the xylem can alter its developmental trajectory.

### Ectopic assembly of the PAX-BRX-PIP5K1 module impacts cellular auxin response

Consistent with the morphological observations, *ARABIDOPSIS HISTIDINE PHOSPHOTRANSFERPROTEIN 6 (AHP6)*, a key promoter of protoxylem formation (Mahonen et al., 2006; Moreira et al., 2013), *INDOLE-3-ACETIC ACID INDUCIBLE 19 (IAA19)*, a xylem-expressed auxin-inducible gene (Muto et al., 2007), and *VASCULAR RELATED NAC-DOMAIN PROTEIN 7 (VND7)*, an auxin-responsive master regulator of xylem differentiation (Yamaguchi et al., 2011; Hirai et al., 2019; von der Mark et al., 2022) were significantly upregulated in plants expressing the entire module as determined by qPCR (Figure 6) (Fig. 6A and B). Such differential expression was not observed with several other genes, including *ARABIDOPSIS THALIANA HOMEOBOX GENE 8 (ATHB8)*, a promoter of procambial cell fate, or *VND6*, a redundant yet not auxin-responsive *VND7* homolog (Kubo et al., 2005; Ramachandran et al., 2021) (Fig. 6C).

**Figure 6.**
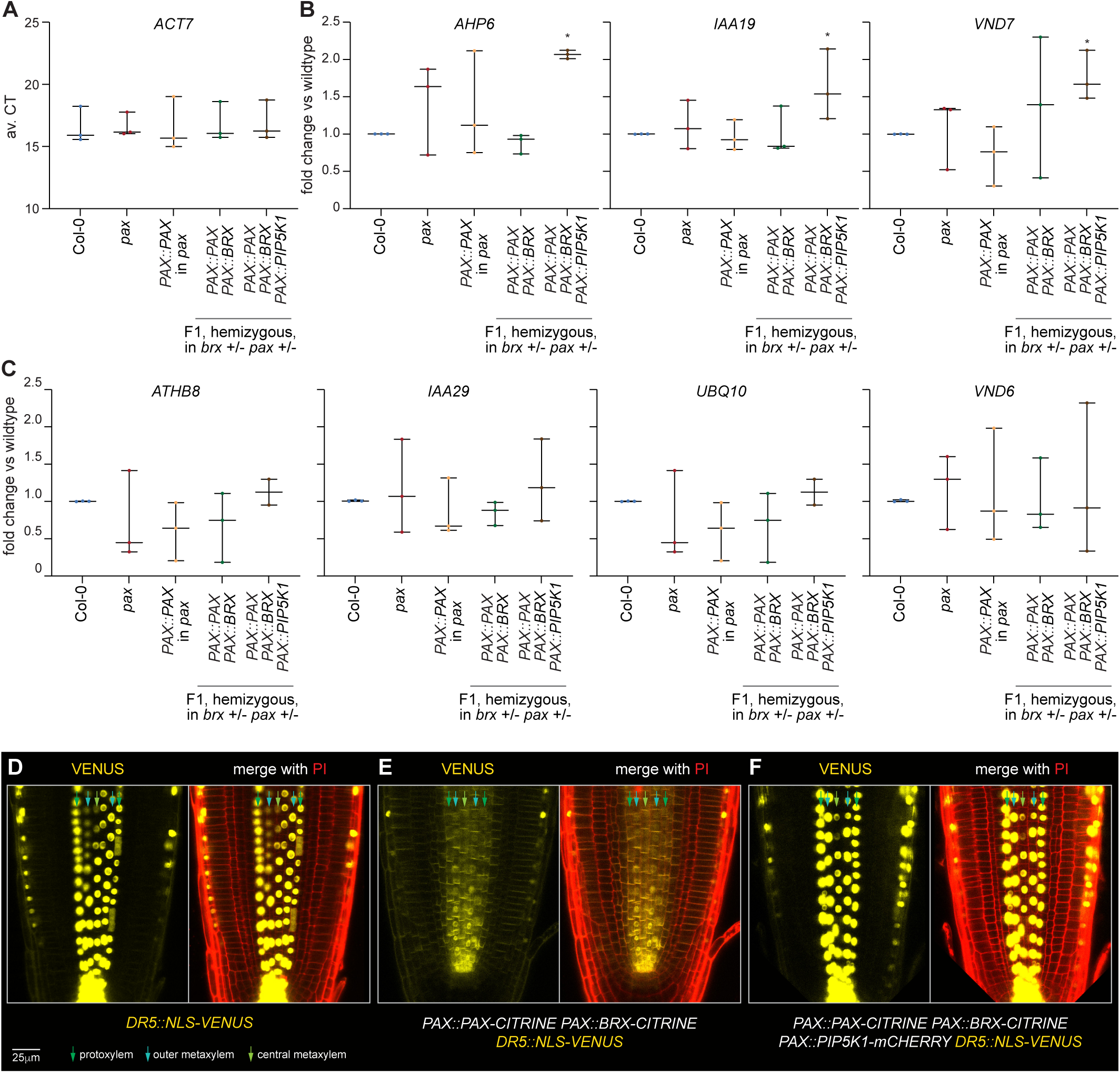
Ectopic PAX-BRX-PIP5K1 assembly affects xylem differentiation markers and auxin activity. (A-C) qPCR quantification of selected xylem development markers and control genes, normalized with respect to expression of the *ACTIN 7 (ACT7)* housekeeping gene (A) and expressed as relative fold-change as compared to Col-0 wildtype (B-C). Plots display the averages of 3 technical replicates from 3 biological replicates each. Statistically significant differences (asterisks) were determined by Student’s t-test compared to Col-0 wildtype, p<0.001 *(AHP6),*, p=0.042 *(IAA19)*, p=0.008 *(VND7).* (D-F) Confocal microscopy images of the auxin activity reporter *DR5::NLS-VENUS* in Col-0 wildtype or in the presence of the indicated transgenes. Yellow fluorescence: NLS-VENUS (nuclear signal) or PAX/BRX-CITRINE (plasma membrane signal); Red fluorescence: propidium iodide (PI) signal. Box plots display 2nd and 3rd quartiles and the median, bars indicate maximum and minimum.

The PAX-BRX-PIP5K module has an important role in guiding the transition of developing PPSEs towards differentiation (Marhava et al., 2020), which has been correlated with timely auxin accumulation (Marhava et al., 2018; Moret et al., 2020; Aliaga Fandino and Hardtke, 2022). In developing PPSEs of *brx* mutants, auxin levels as determined by the DII-VENUS reporter are generally lower and more variable than in wildtype (Marhava et al., 2018), and here we consistently found the same for *pax* mutants (Fig. S6A and B). Moreover, auxin levels were also reduced in developing metaxylem vessels (Fig. S6C and D), which may reflect the systemic impact of perturbed protophloem development (Anne and Hardtke, 2017). To investigate whether a module gain-of-function affects auxin levels, we crossed combinations of our transgenes with the transcriptional *DR5::NLS-VENUS* auxin reporter line (Heisler et al., 2005). These plants displayed the same root phenotypes observed earlier, confirming the dominant effects. Moreover, compared to wildtype, auxin activity was strongly reduced in the developing xylem of PAX + BRX double transgenics (Fig. 6D-E), whereas auxin response appeared to be stronger in PAX + BRX + PIP5K1 triple transgenics (Fig. 6F). These observations show that ectopic assembly of the PAX-BRX-PIP5K1 module in developing xylem vessels alters cellular auxin activity, likely by its impact on trans-cellular auxin flux.

## Discussion

PIN-mediated auxin efflux is subject to complex regulatory inputs, among which targeted PIN recycling and activation are most prominent (Kleine-Vehn et al., 2011; Adamowski and Friml, 2015; Barbosa et al., 2018). AGC kinases play a key role in these processes, through phosphorylation of PIN proteins in their cytoplasmic hydrophilic loop, which is for example necessary for PIN-mediated auxin efflux in the heterologous *Xenopus laevis* oocyte system (Zourelidou et al., 2014; Weller et al., 2017). Several experimentally verified PIN phosphosites have been described and their combinatorial state in a yet to be fully understood ‘phosphocode’ may determine the overall activity and turnover of PINs (Bassukas et al., 2022). Despite their generally close phylogenetic relation and structural similarity (Galvan-Ampudia and Offringa, 2007), AGC kinases have diverged in their effects on PIN activity. For example, although both PID- and D6PK-mediated PIN phosphorylation promotes auxin efflux in the oocyte system (Zourelidou et al., 2014; Weller et al., 2017), *in planta* PID, but not D6PK, also triggers PIN relocalization through transcytosis that competes with basal endocytic PIN recycling (Kleine-Vehn et al., 2009; Dhonukshe et al., 2010; Weller et al., 2017). Similar to PID and D6PK, PAX can stimulate PIN-mediated auxin efflux in the oocyte system, but compared to those other kinases it is a relatively weak activator (Marhava et al., 2018). However, a phosphomimic PAX version that simulates the auxin-stimulated PAX phosphorylation by PDKs is not only a much stronger activator of auxin efflux in the oocyte system, but also hyperactive *in planta* (Marhava et al., 2018; Xiao and Offringa, 2020). Finally, what sets PAX apart from the other kinases in its family is its unique N-terminus (Galvan-Ampudia and Offringa, 2007), which was recently shown to be necessary for interaction with PIP5K (Wang et al., 2023). Here we found that *in planta*, PIP5K recruitment dampens PAX inhibition by BRX as demonstrated by phenotypic read-outs as well as cellular features, notably S231 phosphorylation of PIN1. Our data reiterate that PIP5K promotes PAX activity (Wang et al., 2023), and suggest that S231 phosphorylation of PIN1 by PAX triggers PIN1 endocytosis and subsequent turnover. Moreover, we found that unlike PID, PAX cannot induce PIN depolarization. This may be related to the fact that (ectopically expressed) PID is largely apolar in PPSEs (Wang et al., 2023), whereas PAX remains polar localized even upon prolonged induction. In summary, our results suggest that PAX control of PIN activity is fundamentally distinct from both PID and D6PK due to its unique N-terminus, which allows interaction with PIP5K.

Nevertheless, also in the ectopic xylem context, efficient PIP5K recruitment to the PIN domain requires BRX. Our data thus reiterate that PAX activity depends on its intricate quantitative relationship with both BRX and PIP5K (Marhava et al., 2020; Wang et al., 2023). Until now, the collective, dynamic steady-state output of this three-protein module remained unclear however, because whereas the oocyte assays suggested that PAX kinase activity primarily stimulates PIN-mediated auxin efflux (Marhava et al., 2018; Koh et al., 2021), the PAX-dependent PIN abundance minimum, the ‘donut’ pattern, suggested that PAX kinase activity may also reduce PIN-mediated auxin efflux (Marhava et al., 2020; Wang et al., 2023). The observation that pertinent PIN phosphomimic variants are not functional *in planta* (Huang et al., 2010; Weller et al., 2017) suggests that the two processes could also be intricately linked. Thus, PAX-mediated PIN1 phosphorylation may transiently stimulate auxin efflux but also promote its eventual reset through PIN internalization. This would reconcile a rheostat function that coordinates auxin levels between adjacent cells along a file with a canalization function that nevertheless promotes auxin accumulation in those cells as compared to their lateral neighbors.

Expression of the PAX-BRX-PIP5K1 module in the developing xylem allowed us to monitor the output of these proposed dynamics in an ectopic context in wildtype background and in the absence of CLE45-BAM3 signaling, which interferes with module assembly in developing PPSEs (Diaz-Ardila et al., 2023; Wang et al., 2023). Monitoring of an auxin activity reporter indicates that expression of the PAX-BRX rheostat suppresses auxin accumulation in the developing xylem. Given the systemic importance of xylem-derived auxin for root meristem development (Bishopp et al., 2011), this may explain the short root phenotype and shorter xylem cells of the pertinent transgenic lines. By contrast, ectopic assembly of the entire PAX-BRX-PIP5K module results in higher auxin activity likely due to an overall net auxin retention, which correlates with enhanced PIN patterning (i.e. lower PIN abundance) and an accelerated trajectory of xylem vessel differentiation. These findings are consistent with the relatively lower auxin levels we observed in the xylem of *pax* mutants, and the recent demonstration that xylem vessel differentiation requires auxin accumulation (von der Mark et al., 2022). Thus, in summary our observations suggest that the PAX-BRX-PIP5K1 module promotes cellular auxin retention and thereby promotes the timely differentiation of developing PPSEs (Marhava et al., 2018; Moret et al., 2020). Since this property can be transferred to the ectopic xylem context, our results also support the notion that the renewed increase of cellular auxin generally observed with reporters after the meristematic cell proliferation stage (Santuari et al., 2011; Brunoud et al., 2012) is likely a generic cue for the timing of differentiation across root tissues.

## Materials and Methods

### Plant materials and growth conditions

*A. thaliana* accession Columbia-0 (Col-0) was the wildtype background for all lines used or produced in this study. The following mutant lines and transgenes have been described previously: *brx* (Rodrigues et al., 2009); *pax* and *PAX::PAX-CITRINE* (Marhava et al., 2018); *BRX::BRX-CITRINE* (Rodriguez-Villalon et al., 2014); *CVP2^XVE^::PAX-CITRINE*, *CVP2^XVE^::BRX-CITRINE*, *CVP2^XVE^::PIP5K1-CITRINE*and *CVP2^XVE^::PID-CITRINE* (Wang et al., 2023), *DR5::NLS-VENUS* (Heisler et al., 2005), *35S::mDII-VENUS* and *35S::DII-VENUS* lines in Col-0 and *brx* (Santuari et al., 2011; Brunoud et al., 2012; Marhava et al., 2018).

### Growth conditions

Seeds of *A. thaliana* were surface sterilized and then stratified for 2 days in the dark at 4°C before germination and growth in continuous light at 22°C on vertically placed Petri dishes that contained 0.5× Murashige and Skoog (MS) media supplemented with 0.8% agar and 0.3% sucrose.

### Constructs and generation of transgenic lines

Transgenes for plant transformation were created in suitable binary vectors using standard molecular biology procedures. For the *PAX::BRX-CITRINE* and *PAX::PIP5K1-mCHERRY* constructs, the *PAX* promoter region (Marhava et al., 2018) was amplified and cloned into pDONR P4P1R. The genomic fragments of the *PIP5K1* and *BRX* transcript regions, without their STOP codons, were amplified and cloned into pDONR 221. These entry clones together with CITRINE or 2xmCHERRY in pDONR P2RP3 were combined into the destination vector pH7m34GW by the multisite Gateway recombination system. To generate the inducible *CVP2^XVE^::C-HUB-CITRINE* and *CVP2^XVE^::AUXILINE-LIKE2-CITRINE*fusions, the *CVP2^XVE^* promoter (Wang et al., 2023) region was amplified and cloned into pDONR P4P1R, the *AUXILINE-LIKE2* (At4g12770) (Adamowski et al., 2018) and C-HUB (Dhonukshe et al., 2007) coding sequences without their STOP codons were cloned into pDONR 221 and the CITRINE coding sequence into pDONR P2RP3. These entry clones were combined into binary vector pH7m34GW. The binary constructs were introduced into *Agrobacterium tumefaciens* strain GV3101 pMP90 and transformed into the pertinent Arabidopsis genotypes using the floral dip method. For the *35S::mDII-VENUS* and *35S::DII-VENUS* lines in *pax* mutant, the two constructs in Col-0 background were crossed into *pax* and selected by genotyping. For transgene combinations, *PAX::PAX-CITRINE* in *pax* background and *PAX::BRX-CITRINE* with or without *PAX::PIP5K1-mCHERRY* in *brx* background were crossed to create hemizygous F1 plants.

### Auxin treatments

To monitor auxin response of BRX, 5-day-old seedlings were transferred into liquid MS media with mock or 10μM auxin (1-naphthylacetic acid dissolved in DMSO). Seedlings were removed for analysis after 3h.

### Estradiol treatments

To induce effectors expressed under control of the *CVP2^XVE^* promoter, 5-day-old seedlings were transferred onto plates of ½ MS media supplemented with 5 μM estradiol. Seedlings were removed for analysis at indicated time points.

### Confocal imaging and image processing

Confocal microscopy was performed on *Leica Stellaris 5* and *Zeiss LSM 880 with Airyscan* inverted confocal scanning instruments. To visualize reporter genes and staining signals, the following fluorescence excitation-emission settings were used: CITRINE excitation 514 nm, emission 529 nm; VENUS excitation 515 nm, emission 528 nm; propidium iodide excitation 536 nm, emission 617 nm; Alexa Fluor 488 excitation 498 nm, emission 520 nm; Alexa Fluor 546 excitation 556 nm, emission 573 nm; calcofluor white excitation 405 nm, emission 425–475 nm. Pictures were taken with 20× or 40× water/oil immersion objectives. For presentation, composite images had to be assembled in various instances. Sequential scanning was used for co-localization studies to avoid interference between fluorescence channels. For image analyses, *ImageJ*, *Zeiss Zen 2011 (black edition)*, and *Imaris* image analysis software were used.

### Protein immunolocalization

Whole mount immunolocalization in 5-day-old seedlings was performed as described (Marhava et al., 2020; Wang et al., 2023). Briefly, seedlings were fixed under vacuum in 4 % paraformaldehyde (dissolved in MTSB: 15 gl^−1^ PIPES, 1.9 gl^−1^ EGTA, 1.32 gl^−1^ MgSO4·7 H2O, and 5 gl^−1^ KOH, adjusted to pH 6.8-7.0 with KOH) supplemented with 0.1 % Triton for 50 min. Samples were then washed 3x with MTSB/0.1% Triton and 2x with water for 10 min. For cell wall digestion, samples were treated for 30 min. with 2% driselase in MTSB at 37°C. After washing with MTSB, samples were treated 2x for 30 min. with permeabilization solution (10% DMSO and 3% NP-40 in MTSB). Next, samples were washed 5x with MTSB, pre-incubated in 2% BSA in MTSB for 1 h, and incubated with primary antibody for 4 h at 37°C, then with secondary antibody for 3 h at 37°C. After each antibody treatment, samples were washed 5-7x with MTSB for 10-15 min. Samples were mounted in Citi-fluor antifade mounting medium and imaged by confocal laser-scanning microscopy. Separation of individual cells, if desired, was achieved by applying light thumb pressure on slides before imaging. The primary antibody dilutions were: 1:500 for anti-GFP mouse; 1:600 for anti-GFP rabbit; 1:500 for anti-BRX rabbit; 1:250 for anti-PIN1 goat; 1:100 for anti-PIN1 J231 rabbit; 1:300 for anti-PIN1 J271 rabbit; 1:500 for anti-PAX rabbit. The secondary antibody dilutions were: 1:500 for Alexa Fluor 488 anti-mouse; 1:500 for Alexa Fluor 546 anti-rabbit; 1:500 for Alexa Fluor 546 anti-goat.

### qPCR

For expression analysis, ca. 7mm of the root tip from 7-day-old seedlings of each genotype were collected. Total RNA was extracted using an RNeasy Plant Mini Kit (QIAGEN) and treated with DNase I on the column. cDNA was synthesized with the SuperScriptII kit (Invitrogen) and used as a template for qPCR assays with the MESA BLUE kit (Takyon), using the primers listed in Supplemental Table S1. The relative expression values were calculated using the *ACTIN 7* gene as a reference, using the ΔΔCT method. All assays were performed with three technical replicates each of three biological replicates.

### Xylem differentiation quantification

To analyze xylem differentiation status, roots were mounted in chloralhydrate solution (8:2:1 chloralhydrate:glycerol:water w/v/v), and visualized on a *Leica* light microscope with differential interference contrast optics. To score trajectories in the meristem, 6-day-old plants were fixed in 4% PFA, washed 4x in MTSB, and cleared overnight in ClearSee solution. The next day samples were placed in a basic fuchsin-ClearSee mixture (final basic fuchsin concentration 0.2% in ClearSee) overnight. The following day the samples were washed 2x for 1 h in ClearSee and mounted on slides with ClearSee for visualization on the *Leica Thunder* microscope. To count protoxylem precursors from the QC to the first differentiated protoxylem vessel, samples from anti-PIN1 immunolocalization were used to distinguish cell boundaries.

### Statistical analyses

Analyses to determine statistical significance were performed in Graphpad Prism software, version 9.3.1. Specific statistical tests used (Student’s t-test, Fisher’s exact test, ordinary one-way ANOVA) are indicated in the figure legends and were always two-tailed. Robust regression and outlier removal (ROUT) analyses were performed on discrete measurements to detect (rare) outliers, which were removed. All experiments were replicated at least twice, typically three times.

## Data availability

This study includes no data deposited in external repositories.

## Acknowledgments

We would like to thank Prof. N. Geldner for comments on the manuscript, Prof. C. Schwechheimer for a gift of phosphosite-specific anti-PIN1 antibodies, and Prof. R. Offringa for PDK-related materials. This study was funded by a bilateral grant between the *Swiss National Science Foundation* (SNF) and the *Czech Science Foundation* (CSF) (SNF grant 310030L_197794 awarded to C.S.H. and CSF grant 21-08021L awarded to J.P.). A.C.A.F. was supported by a Ph.D. Fellowship from the Faculty of Biology and Medicine of the University of Lausanne.

## Author contributions

Conceptualization A.C.A.F and C.S.H.; Methodology A.C.A.F., A.J. and P.M.; Investigation A.C.A.F., A.J. and P.M.; Validation A.C.A.F., A.J. and P.M.; Visualization A.C.A.F., A.J. and P.M.; Writing – Original Draft A.C.A.F. and C.S.H.; Writing – Review & Editing A.C.A.F., A.J., P.M., J.P. and C.S.H.; Funding Acquisition J.P. and C.S.H.; Supervision J.P. and C.S.H.

## Disclosure and competing interests statement

The authors declare no competing interests.

---

**Figure S1.**
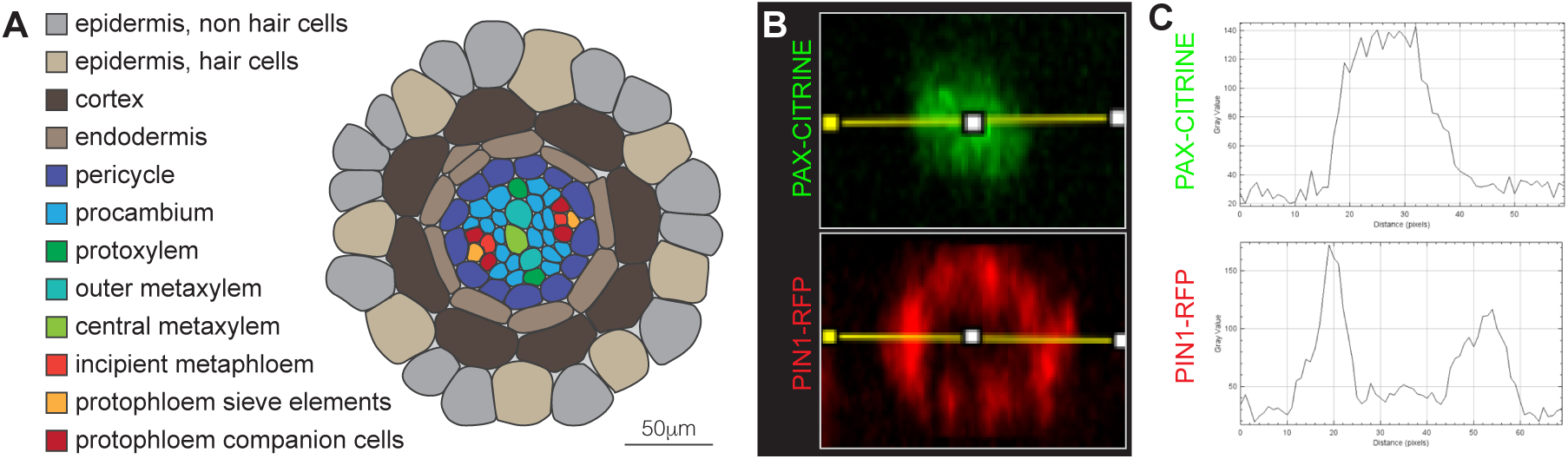
Root tissue layers and subcellular PIN pattern. (A) Schematic presentation of an Arabidopsis primary root cross-section at the level of PPSE differentiation. (B) Confocal microscopy live images of PAX-CITRINE and PIN1-RFP fusion proteins at the rootward plasma membrane of a developing PPSE, illustrating the central ‘muffin’ localization of PAX that is complementary to the ‘donut’ pattern of PIN1. (C) Signal intensity traces across the images shown in (B).

**Figure S2.**
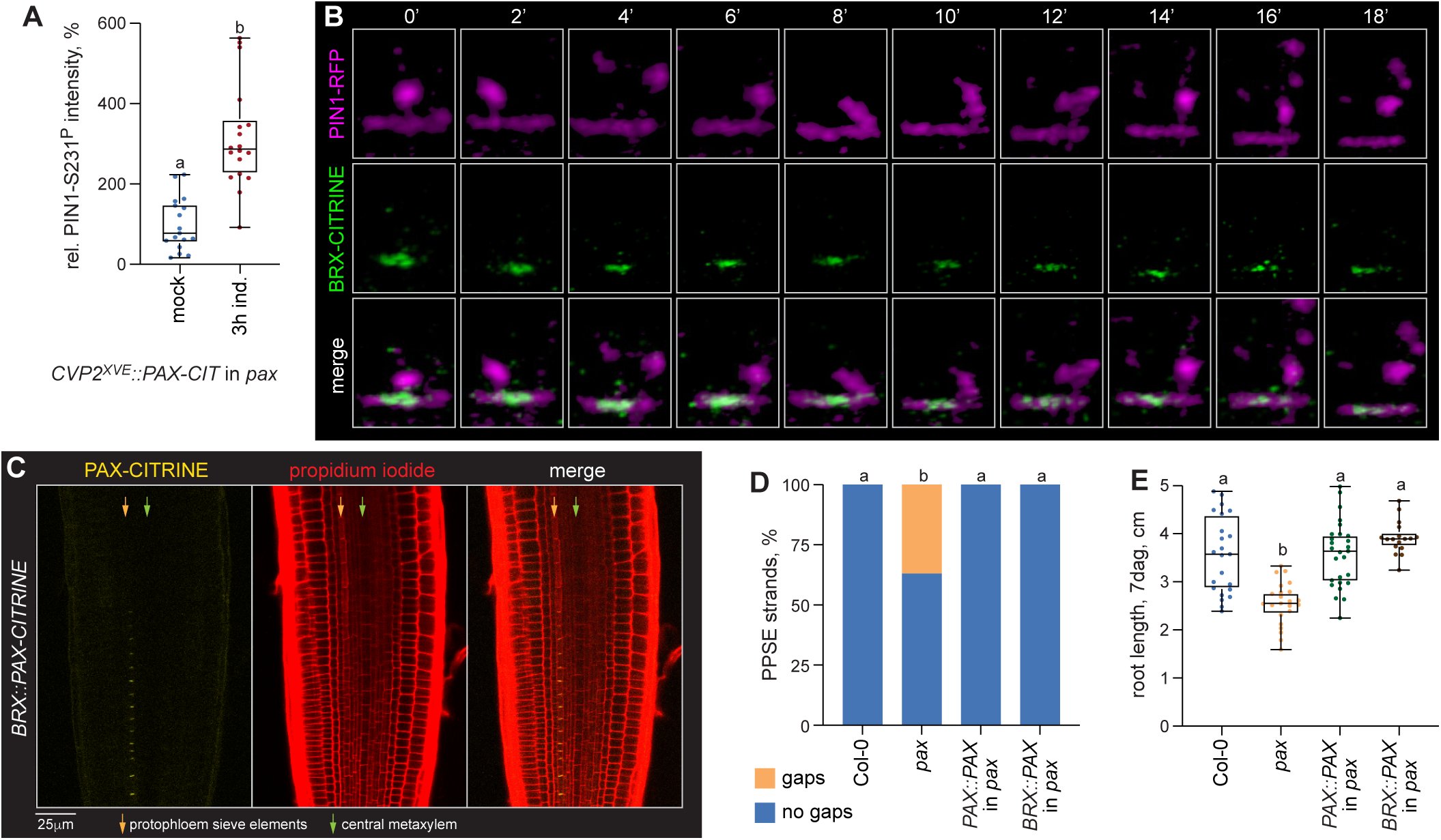
PIN1 endocytosis and PAX activity in developing PPSEs. (A) Relative signal intensity of S23l^p^-specific PIN1 immunostaining in developing PPSEs upon estradiol-induction of PAX-CITRINE fusion protein under control of the *CVP2^XVE^* promoter. n=17-18 roots; statistically significant difference (lower case letters) was determined by Student’s t-test, p<0.0001. (B) Time course of PIN1-RFP (magenta fluorescence) and BRX-CITRINE (green fluorescence) fusion protein dynamics at the rootward plasma membrane of a developing PPSE, capturing PIN1-RFP internalization from the center. (C) Confocal live imaging of PAX-CITRINE fusion protein (yellow fluorescence, left panel) expressed under control of the *BRX* promoter in *pox* mutant background, and propidium iodide cell wall staining (red fluorescence, center panel). (D) Scoring of PPSE differentiation failures (’gaps’) in the indicated genotypes (PAX was expressed as a CITRINE fusion protein). n=30-52 PPSE cell files; statistically significant differences (lower case letters) were determined by ordinary one-way ANOVA, p<0.0001. (E) Primary root length of indicated genotypes (PAX was expressed as a CITRINE fusion protein). n=17-27 roots; statistically significant differences (lower case letters) were determined by ordinary one-way ANOVA, p<0.0001. Box plots display 2nd and 3rd quartiles and the median, bars indicate maximum and minimum.

**Figure S3.**
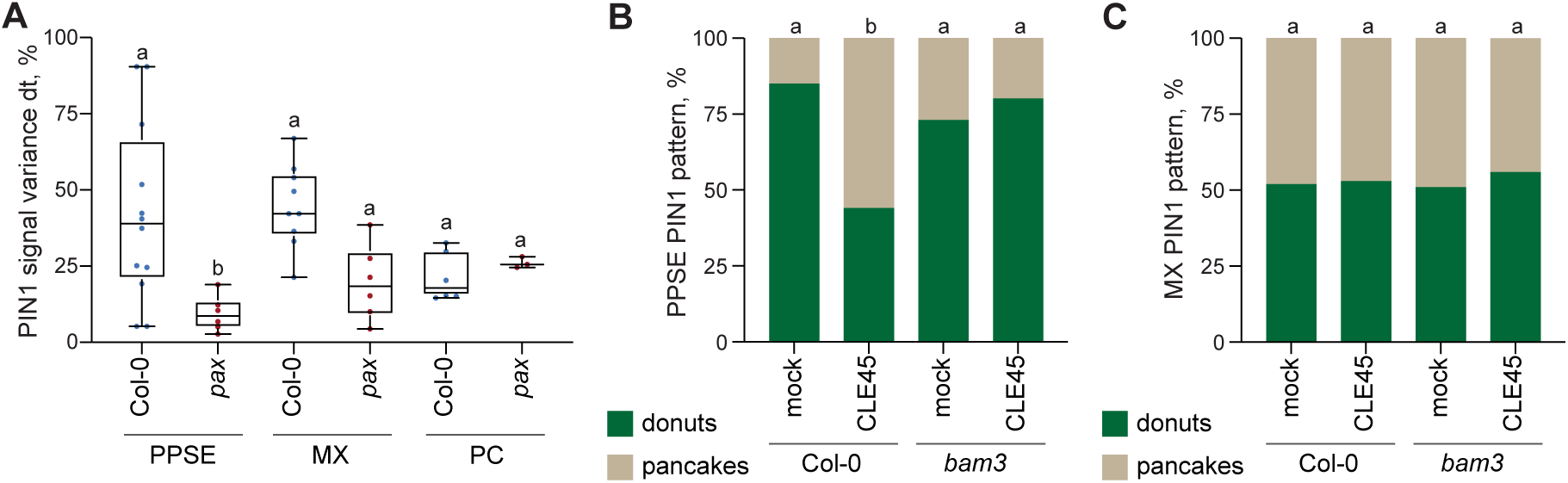
PIN1 dynamics and CLE45 response. (A) PIN1-RFP fusion protein dynamics in different tissues as determined by the signal intensity variance over a 10 min. time course (PC: procambial cells). n=3-12 cells; statistically significant differences (lowercase letters) compared to Col-0 wildtype were determined by ordinary one-way ANOVA, p<0.0137. (B-C) Quantification of the subcellular PIN1 pattern in developing PPSEs (B) or MX vessels (C) in Col-0 wildtype or *bom3* mutant background, in mock conditions or upon 3h treatment with 10 micromolar CLE45 peptide. n=52-106 PPSEs (B) and n=94-153 MX vessels (C); statistically significant differences (lower case letters) were determined by Fisher’s exact test, p<0.001. Box plots display 2nd and 3rd quartiles and the median, bars indicate maximum and minimum.

**Figure S4.**
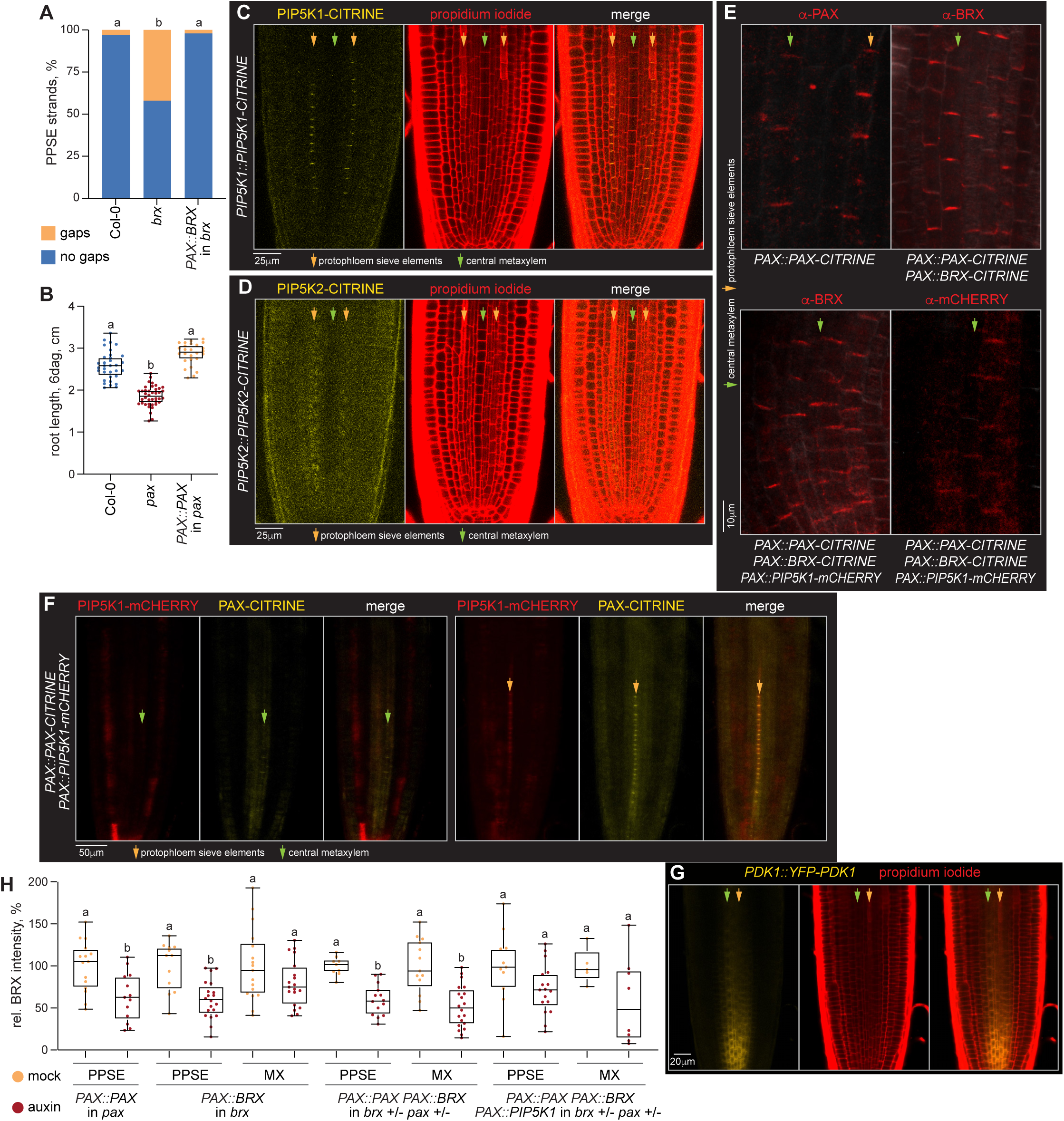
Transgene expression patterns. (A) Scoring of PPSE differentiation failures (’gaps’) in the indicated genotypes (BRX was expressed as a CITRINE fusion protein). n=22-58 PPSE cell files; statistically significant differences (lower case letters) were determined by Fisher’s exact test, p<0.001. (B) Primary root length of indicated genotypes (PAX was expressed as a CITRINE fusion protein). n=26-40 roots; statistically significant differences (lower case letters) were determined by ordinary one-way ANOVA, p<0.0001. (C-D) Confocal live imaging of PIP5K1 and PIP5K2 CITRINE fusion proteins (yellow fluorescence, left panels) expressed under control of their native promoters in *pip5k2* mutant background, and corresponding propidium iodide cell wall staining (red fluorescence, center panels). (E) Detection of PAX, BRX and PIP5K1 CITRINE/mCHERRY fusion proteins by immunostaining (red fluorescence) with indicated antibodies in the indicated transgenic genotypes, illustrating (ectopic) expression in the xylem axis and polar plasma membrane association. (F) Confocal live imaging of PAX-CITRINE and PIP5Kl-mCHERRY fusion proteins in wildtype background. (G) Confocal live imaging of YFP-PDK1 fusion protein (yellow fluorescence, left panel) expressed under control of its native promoter in *pdkl pdk2* double mutant background, and propidium iodide cell wall staining (red fluorescence, center panel). (H) Quantification of auxin-induced (10 micromolar auxin treatment, 3h) plasma-membrane-dissociation of BRX-CITRINE fusion protein in developing PPSEs and MX vessels in the presence of indicated transgene combinations (relative average signal intensities per root). n=5-20 roots; statistically significant differences (lower case letters) compared to mock were determined by ordinary one-way ANOVA, p≤0.0313. Box plots display 2nd and 3rd quartiles and the median, bars indicate maximum and minimum.

**Figure S5.**
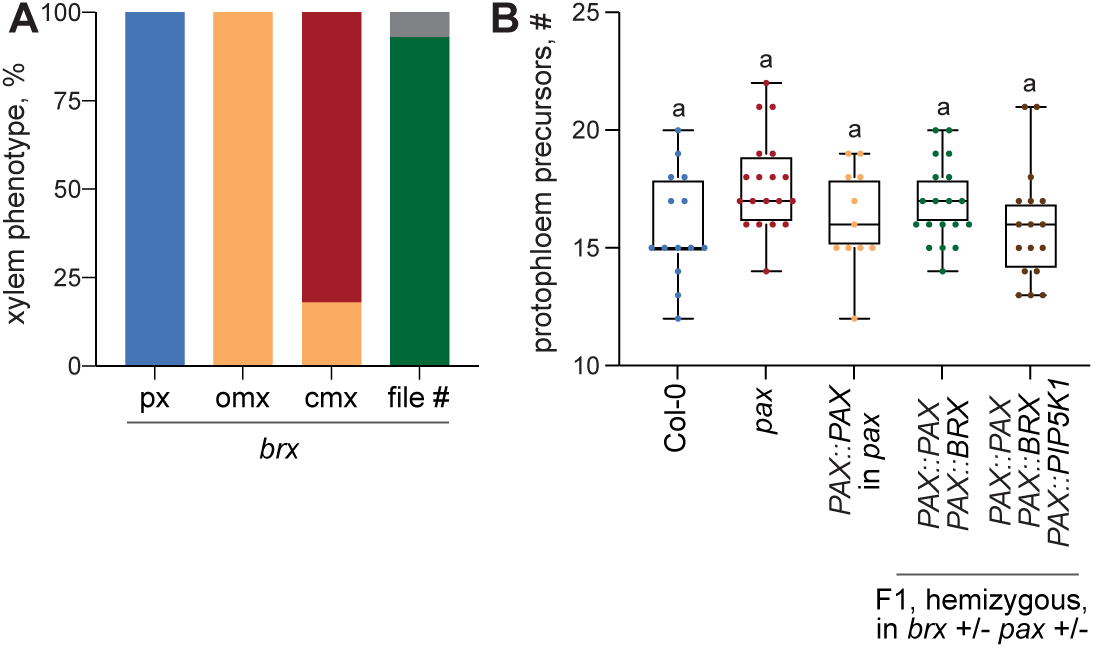
Xylem differentiation in *brx,* and PPSE precursors. (A) Quantification of differentiation status per vessel type in *brx* mutant background, determined by differential interference contrast light microscopy 5-7 mm above the root tip. n=28 roots. (B) Number of undifferentiated precursors in PPSE cell files of the indicated genotypes, counted from the QC. n=ll-19 cell files; statistically significant differences (lower case letters) were determined by ordinary one-way ANOVA. Box plots display 2nd and 3rd quartiles and the median, bars indicate maximum and minimum.

**Figure S6.**
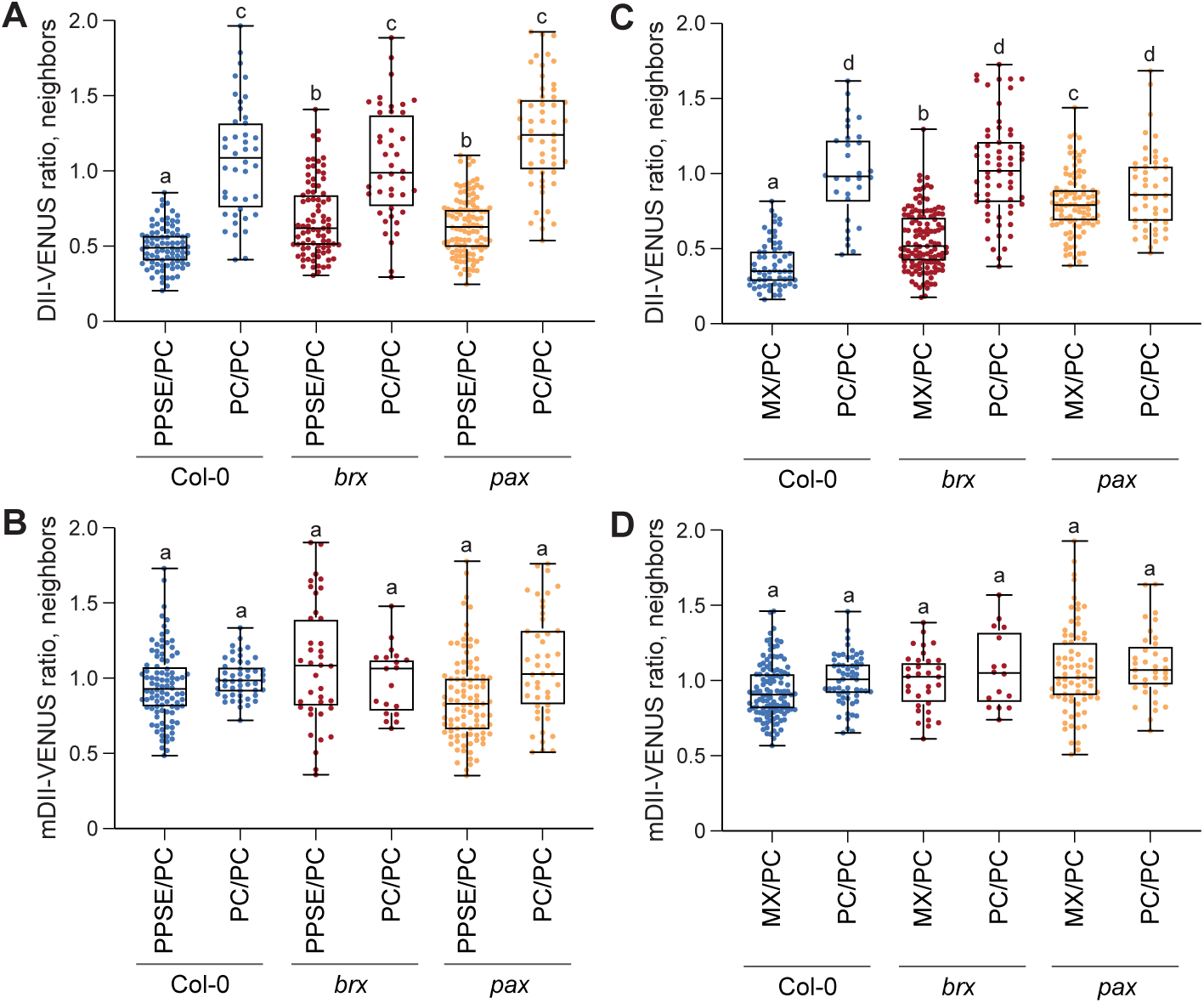
Auxin reporter quantification. (A-D) Quantification of the auxin-degradable Dll-VENUS reporter and the auxin-resistant mDII-VENUS reporter in different tissues of Col-0 wildtype or *pax* or *brx* mutants. Signal intensities were determined by confocal live imaging to calculate the signal intensity ratio between adjacent cells in neighboring cell files. n=16-121 cell pairs; statistically significant differences (lower case letters) were determined by ordinary one-way ANOVA, p≤0.0007. Box plots display 2nd and 3rd quartiles and the median, bars indicate maximum and minimum.

## Notes

### Competing Interest Statement

The authors have declared no competing interest.

